# Predicting the effects of deep brain stimulation using a reduced coupled oscillator model

**DOI:** 10.1101/448290

**Authors:** Gihan Weerasinghe, Benoit Duchet, Hayriye Cagnan, Peter Brown, Christian Bick, Rafal Bogacz

**Affiliations:** MRC Brain Network Dynamics Unit, Nuffield Department of Clinical Neurosciences, University of Oxford, Oxford, UK.; Oxford Centre for Industrial and Applied Mathematics, Mathematical Institute, University of Oxford, Oxford, UK.; Centre for Systems Dynamics and Control and Department of Mathematics, University of Exeter, Exeter, UK.

## Abstract

Deep brain stimulation (DBS) is known to be an effective treatment for a variety of neurological disorders, including Parkinson’s disease and essential tremor (ET). At present, it involves administering a train of pulses with constant frequency via electrodes implanted into the brain. New ‘closed-loop’ approaches involve delivering stimulation according to the ongoing symptoms or brain activity and have the potential to provide improvements in terms of efficiency, efficacy and reduction of side effects. The success of closed-loop DBS depends on being able to devise a stimulation strategy that minimizes oscillations in neural activity associated with symptoms of motor disorders. A useful stepping stone towards this is to construct a mathematical model, which can describe how the brain oscillations should change when stimulation is applied at a particular state of the system. Our work focuses on the use of coupled oscillators to represent neurons in areas generating pathological oscillations. Using a reduced form of the Kuramoto model, we analyse how a patient should respond to stimulation when neural oscillations have a given phase and amplitude. We predict that, provided certain conditions are satisfied, the best stimulation strategy should be phase specific but also that stimulation should have a greater effect if applied when the amplitude of brain oscillations is lower. We compare this surprising prediction with data obtained from ET patients. In light of our predictions, we also propose a new hybrid strategy which effectively combines two of the strategies found in the literature, namely phase-locked and adaptive DBS.

**Author summary:** Deep brain stimulation (DBS) involves delivering electrical impulses to target sites within the brain and is a proven therapy for a variety of neurological disorders. Closed loop DBS is a promising new approach where stimulation is applied according to the state of a patient. Crucial to the success of this approach is being able to predict how a patient should respond to stimulation. Our work focusses on DBS as applied to patients with essential tremor (ET). On the basis of a theoretical model, which describes neurons as oscillators that respond to stimulation and have a certain tendency to synchronize, we provide predictions for how a patient should respond when stimulation is applied at a particular phase and amplitude of the ongoing tremor oscillations. Previous experimental studies of closed loop DBS provided stimulation either on the basis of ongoing phase or amplitude of pathological oscillations. Our study suggests how both of these measurements can be used to control stimulation. As part of this work, we also look for evidence for our theories in experimental data and find our predictions to be satisfied in one patient. The insights obtained from this work should lead to a better understanding of how to optimise closed loop DBS strategies.

## Introduction

Symptoms of several neurological disorders are thought to arise from overly synchronous activity within neural populations. The severity of clinical impairment in Parkinson’s disease (PD) is known to be correlated with an increase in the beta (13-35 Hz) oscillations in the local field potential (LFP) and in the activity of individual neurons in the basal ganglia [1, 2]. The tremor symptoms associated with essential tremor (ET) are thought to arise from synchronous activity in a network of brain areas including the thalamus [3]. In both PD and ET the muscle activity driving the tremor is coherent with local field potentials in the thalamus [4] and bursts of spikes produced by individual thalamic neurons [5].

Deep brain stimulation (DBS) is a well-established treatment option for PD and ET which involves delivering stimulation via electrodes implanted into the brain. The present generation of the technology involves manually tuning the parameters of stimulation, such as the pulse width, frequency and intensity in an attempt to achieve the best treatment. In particular, the choice of frequency is known to be crucial for efficacy, and high frequency DBS (120-180 Hz) has been found to be effective for PD and ET patients [6]. High frequency DBS is known to suppress the pathological oscillations occurring in PD [7], but despite its long history, the underlying mechanisms causing this suppression remain unclear, and several distinct theories have been proposed [8–10]. One influential theory suggests that high frequency DBS activates target neurons to such an extent that their synaptic transmission becomes saturated and they are no longer able to transmit pathological oscillations [11]. Since high frequency DBS can cause side-effects such as speech-impairments [12] and gambling tendencies [13], improvements to this treatment approach are desirable.

It is thought that improvements could be achieved if future devices were to operate ‘closed-loop’, delivering stimulation only when needed and according to the ongoing symptoms of the patient [14]. A number of approaches to closed-loop DBS can be found in the literature and of these we focus on two, namely adaptive DBS [15] and phase-locked DBS [16]. In adaptive DBS high frequency stimulation is applied only when the amplitude of oscillations exceed a certain threshold [15]. In phase-locked DBS stimulation is applied according to the instantaneous phase of the oscillations, which for ET patients corresponds to stimulation at roughly the tremor frequency (typically ~ 5 Hz) [16]. The principles behind both of these approaches are illustrated in Figure 1. Together, these studies suggest that the effects of DBS are dependent on both the phase and amplitude of the oscillations at the time of stimulation.

Modelling the effects of DBS generally poses a challenge since the brain networks involved in disorders such as ET (cortico-thalamic circuit) and PD (cortico-basal-gangla circuit) are complex and it is still debated from which parts of these circuits the pathological oscillations originate [17, 18]. The task can be made more tractable by considering a simple phenomenological model which does not attempt to explicitly describe the underlying circuits, but rather focuses on general mechanisms leading to the synchronization of neurons. One example of this is the Kuramoto model, [19] where the dynamics of neurons are described using a system of homogeneously coupled oscillators, whose phases evolve according to a set of underlying differential equations. Such models are particularly attractive due to their simplicity and explicit dependence on phase, which makes them convenient for describing the effects of phase-locked stimulation.

Coupled oscillator models have been used before to describe the effects of applying DBS at particular phases of ongoing oscillations [20, 21]. In particular, Wilson and Moehlis [21] have described how the optimal intensity of stimulation should depend on the phase of the ongoing oscillations. The phase-dependent effects of stimulation predicted using the coupled-oscillator model [21] have been shown to generalize to other models, as they have been also observed in a biologically realistic model of a neural circuit generating oscillations related to PD [22]. However, to the best of our knowledge, the predictions of these models have not been directly compared with the experimental effects of closed-loop DBS. Furthermore, it has not been described how the effects of DBS should depend on another important characteristic of the ongoing oscillations-namely, the amplitude. In this paper, we attempt to understand the mechanisms which may give rise to the effects of phase-locked DBS by using a reduced Kuramoto model. We analyse how the effects of stimulation should depend on the phase and amplitude of the ongoing tremor oscillations. In addition to this, we compare our predictions with previously obtained experimental data.

## Models

### Neural oscillators

We aim to describe how an underlying system of neural oscillators can give rise to oscillations, such as those found in LFP and tremor. In classic coupled oscillator models, neural oscillators correspond to individual neurons which spontaneously produce spikes [23]. In mathematical models of neurons (e.g. the Hodgkin-Huxley model) a state of a neurons is described by a set of variables, and for certain parameters, it produces spikes at regular intervals, thus its variables have periodic behaviour. A description of such a neuron can be simplified by projecting the state of the neuron on its phase, such that the state of each neuron is simply described by a one-dimensional phase variable, which in absence of external input increases with constant rate from phase 0 to 2n, corresponding to an evolution of a neuron from spike to spike [24].

**Fig.1.**
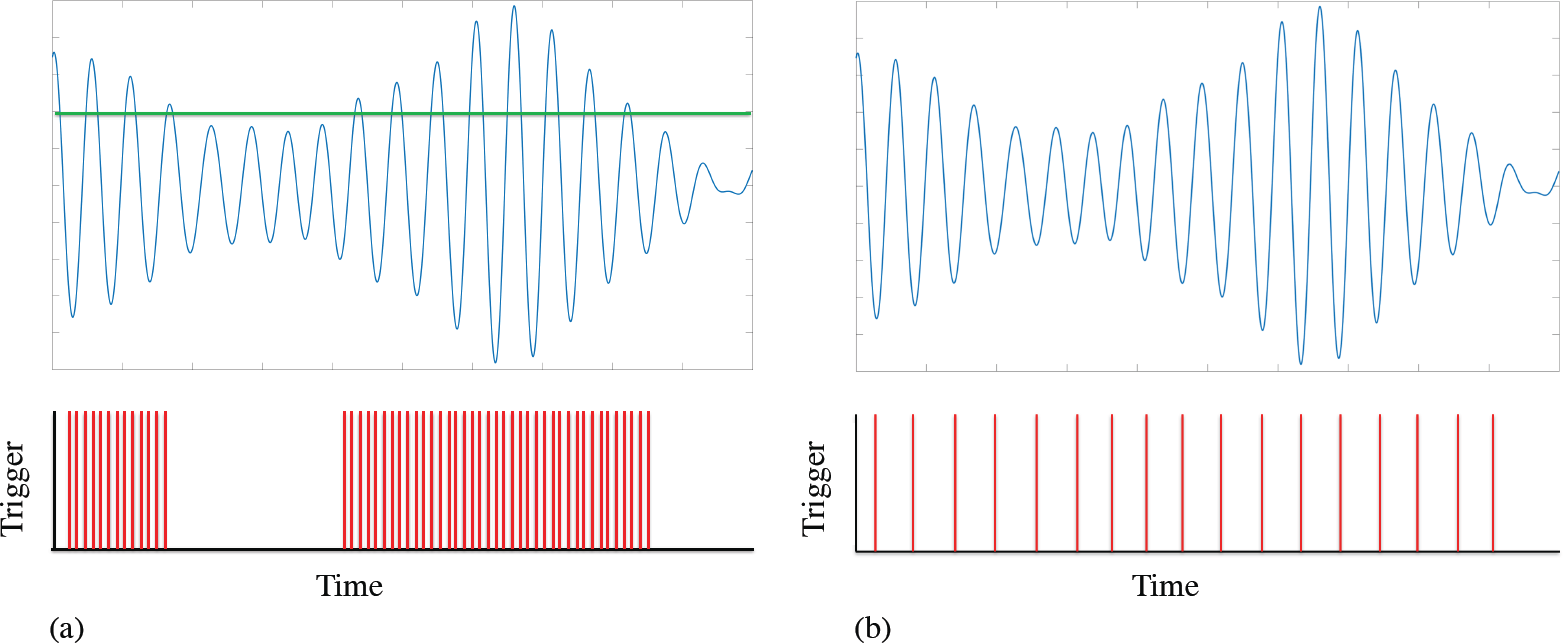
Strategies for closed-loop DBS. (a) Adaptive DBS is delivered only when the amplitude of oscillations exceed a predefined threshold, indicated here by a green line. (b) Phase-locked DBS is delivered only at certain phases of pathological oscillations. In this example, it the stimulation coincides with the troughs of the oscillation.

The above interpretation of neural oscillators as regular-spiking neurons may be suitable for describing relatively fast beta oscillations occurring in PD, but during slower tremor oscillations thalamic neurons produce a burst of activity during a single cycle [5]. Therefore, if one interprets neural oscillators in ET as neurons, the phase would rather describe the changes in a variable governing the burst cycle (e.g. calcium level inside neuron). Alternatively, neural oscillators in ET can be interpreted as micro-circuits, which due to their internal connectivity produce oscillations in the activity of constituent neurons [25]. Since we aim here to develop a general theory, we simply consider the set of *N* neural oscillators with phases {*θ*_m_(t)}. Nevertheless, we make two assumptions about the oscillators related to how they react to input, and how their phases determine the activity of the whole population. In the remainder of this subsection we describe these assumptions and discuss how they could be justified under different interpretations of individual oscillators mentioned above (regular-spiking neurons, bursting neurons and micro-circuits).

First, we assume that if the stimulating input is provided to an oscillator when it is in phase *θ*, its phase will change according to a phase response function *Z*(*θ*). Biological regular-spiking neurons respond more to stimulation when they are already outside their refractory period and their membrane potential is closer to the spiking threshold [26]. Under such conditions, stimulation accelerates a neuron towards spiking. The function *Z*(*θ*) should therefore have higher values in the second part of the spiking cycle. In addition to this and under certain conditions, stimulation just after spiking and during the refractory period can slow a neuron’s spiking [26]. Therefore, *Z*(*θ*) should have negative values for *θ* just above *ψ*. Both of these characteristics can be captured using the phase response function *Z*(*θ*) = — sin(*θ*). We use this simplified phase response function in the first part of our paper and then later consider a more general form. We are not aware of any experimental studies of phase-response curves of bursting neurons, but one could expect that the change in the onset of the next burst will also depend on when during the bursting cycle the stimulation is provided. Phase response functions have been studied in a mathematical model of a micro-circuit composed of connected populations of excitatory and inhibitory neurons [27]. When such micro-circuit receives an input, the phase of the oscillations it produces either advances or reduces depending on when within the cycle the input is provided [27].

Second, we define the average activity of neural oscillators as a superposition of cosine functions, i.e.

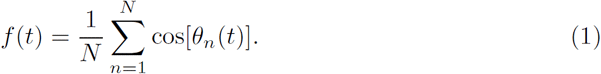

We chose to transform the phase through a cosine function, because this periodic function has a maximum at 0, and in classic coupled oscillator model, phase 0 corresponds to the phase when neurons produce spikes [23]. When regular-spiking neurons are considered, their activity features spikes, rather than varying smoothly like the cosine function, but nevertheless, the effects of spikes on downstream neurons are prolonged in time due to non-instantaneous decay of post-synaptic potentials, thus the cosine function could be seen as a qualitative approximation of the effect of neuron’s activity on downstream cells. If neural oscillators are assumed to correspond to bursting neurons or micro-circuits, the choice of the cosine function is more natural, because their rate of producing spikes varies more gradually. We would assume that the activity of bursting neurons or micro-circuits is highest at phase 0, so the function *f*(*t*) qualitatively reflects the fluctuations in firing rate.

### Relating neural oscillator model to experimental data

The purpose of this subsection is to relate the phases of individual oscillators to the quantities that can be measured from experimental LFP or tremor data, such as the instantaneous phase and the amplitude of the signal.

The function *f*(*t*) is an abstract representation of neural activity and is not directly measurable in typical studies with patients. The experimentally measured signal f_e_(t) is known to be highly correlated to neural activity [5], so it is reasonable to assume f_e_(t) to be some transformed version of *f*(*t*). We assume this transformation to be a simple scaling and shifting, namely

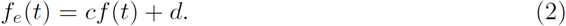

It is common to subtract the mean from a signal, resulting in a signal 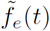 which is independent of *d*. This yields a simple relationship between the experimentally measured data and neural activity,

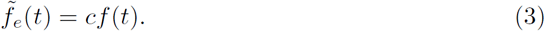

The experimental signal 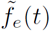 is typically analysed using the Hilbert transform 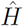 which provides for each time point *t* the values of instantaneous phase and the envelope amplitude *ψ*_e_ and the envelope amplitude *ρ*_e_

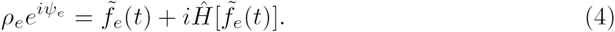

By inserting Eq. (3) into Eq. (4), we can relate the neural activity resulting from coupled oscillators to the experimental phase and amplitude

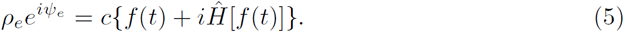

We would like to now relate the experimental phase and amplitude *ρ*_e_ to quantities obtained using the coupled oscillator model. The order parameter for a system of oscillators is defined to be

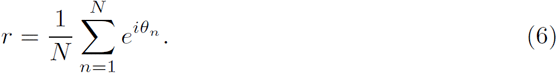

Since *r* is a complex number, it can be written as

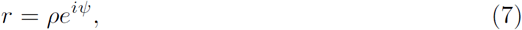

The amplitude *ρ* is a measure of synchrony, with complete desynchrony and synchrony corresponding to 0 and 1, respectively. Using the Euler relation, the order parameter can be written as

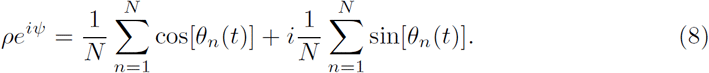

We expect the phases {*θ*_n_} to increase monotonically with time and under these conditions, 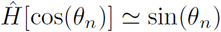. Using this and the expression for the time series given by Eq. (1), Eq.(8) can be written as

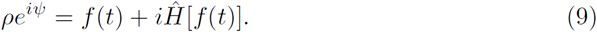

Comparing this with Eq. (5), it can be seen that

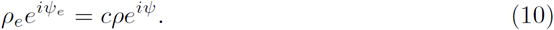

Therefore the experimental amplitude and phase is relatable to the magnitude and phase of the order parameter using

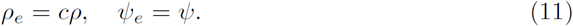

In summary, assuming the experimental data and neural activity are related according to Eq. (2) and that the phases {*θ*_n_} increase monotonically with time, we can use the Hilbert transform of the experimental data to relate the envelope amplitude and instantaneous phase to the magnitude and phase of the order parameter, respectively.

### Kuramoto model

The oscillation data we are concerned with arises from the correlated electrical activity of neural populations. In order to describe such systems we use a coupled oscillator model where the time-evolution for the set of *N* oscillators are given by the Kuramoto equations, with an additional term describing the effects of stimulation [19, 20]

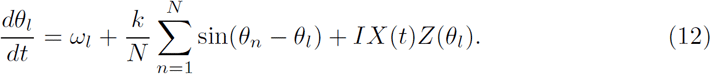

The first term, *ω*_l_ is the natural frequency of oscillator *l*, which describes the frequency in the absence of external inputs. It corresponds to the frequency with which a neuron spontaneously produces spikes or bursts (depending of the interpretation of oscillators introduced above). The second term describes the interactions between oscillators, where k is the coupling constant which controls the strength of coupling between each pair of oscillators and hence their tendency to synchronize. The third term describes the effect of stimulation. The intensity of stimulation is denoted by I and *X*(*t*) is a function which equals 1 if stimulation is applied at time *t* and 0 otherwise. The phase response function for a single oscillator is given denoted by *Z*(*θ*_l_). To avoid confusion, we use ‘intensity’ to refer to the magnitude of stimulation *I*, while the word ‘amplitude’ is used to refer to the amplitude of order parameter *ρ* or of the experimental signal *ρ*_e_. Using the definition of the order parameter given in Eq. (7), Eq. (12) can be transformed to give

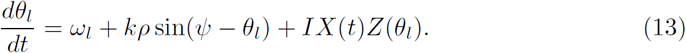

In this form, it is clear that each oscillator has a tendency to move towards the population phase *ψ* and that the strength of this tendency is controlled by the coupling parameter *k*. To gain an intuition for this behaviour readers may wish to explore an online simulation of the model [28].

### Reduced model

In the previous section, we outlined the conditions for which a relationship should hold between the experimental envelope amplitude and instantaneous phase and the quantities associated with the coupled oscillator model, namely the magnitude and phase of the order parameter. A subject’s response to stimulation can be quantified using the amplitude response curve (ARC) and the phase response curve (PRC), which respectively describe changes in the envelope amplitude and phase of the oscillations at the time of stimulation. From a theoretical perspective, the response curves arise from changes to an underlying state of oscillators which in turn gives rise to measurable changes in the ‘macroscopic’ quantities, namely the amplitude and phase of the order parameter. Therefore, in order to obtain an analytical expression for the response curves, we need to know how the order parameter evolves as a function of time. Ott and Antonsen showed that such an expression can be found under the assumption of an infinite system of oscillators and where the distribution of frequencies *g(ω)* is Cauchy with centre *ω*_0_ and width γ. In this section, we summarize their results.

For an infinite system of oscillators, the order parameter can be expressed in terms of the distribution of oscillators *f*(*ω*, *θ*, *t*)

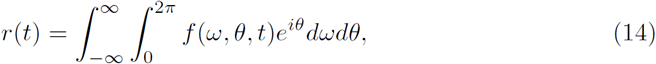

and the time evolution of *f* (*ω*, *θ*, *t*) for the Kuramoto system given by Eq. (13) is given by the continuity equation

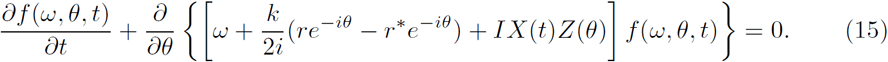

Central to the work of Ott and Antonsen [29] is the use of a guess, or *ansatz,* for the distribution of oscillators given by

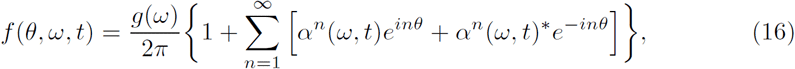

where *α(ω,t*) is a certain function. Ott and Antonsen [29] considered the case of a Kuramoto system with a periodic driving term of strength I and frequency *θ*, whose dynamical equations are given by ^1^

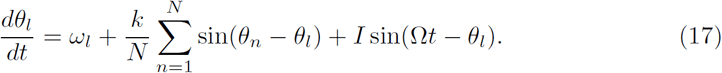

Using this, the *ansatz* and the result that *r(t)* = *α*\*(*w*_0_ — *iγ*,*t*) [30], the time evolution for the order parameter is found to satisfy (for γ = 1)

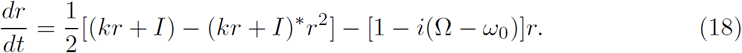

Therefore the state of the Kuramoto system for a large number of oscillators has been reduced from one being described by the set of *N* phases {**θ*_j_*} to one being described by *ρ* and *ψ*.

## Results

### Simplified Response Curves

A patient’s response to phase-locked DBS is typically quantified using the ARC, which describes changes in the envelope amplitude of pathological oscillations (e.g. tremor) as a function of the phase at which the stimulation was delivered. Some studies also report the PRC, which describes changes in the phase of the pathological oscillation as a function of the stimulation phase. Although the effect of phase-locked DBS may also depend on the the amplitude of the ongoing pathological oscillations, this dependence on amplitude has not been analysed before, due primarily to the difficulties associated with obtaining a function of two independent variables from noisy data. Instead, the averaged response curves have been reported [16, 31], which are only functions of the phase and are averaged over the amplitude. Such curves are readily obtainable using standard signal processing techniques.

In this subsection, we derive expressions for the ARC and PRC, where the phase response function for a single oscillator is taken to be *Z*(*θ*) = — sin(*θ*). By setting Ω → 0 and for general γ, Eq. (17) describes a system experiencing an impulse given by the phase response function *Z*(*θ*) = — sin(*θ*).

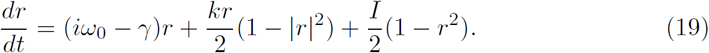

Inserting the expression for the order parameter Eq. (7) into Eq. (19) gives expressions for the time evolution of *ρ* and *ψ*

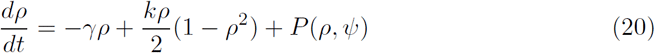

and

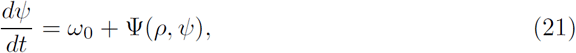

where

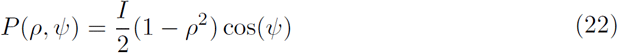

and

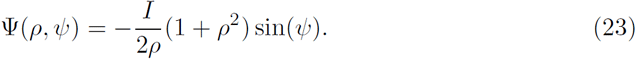

**Fig.2.**
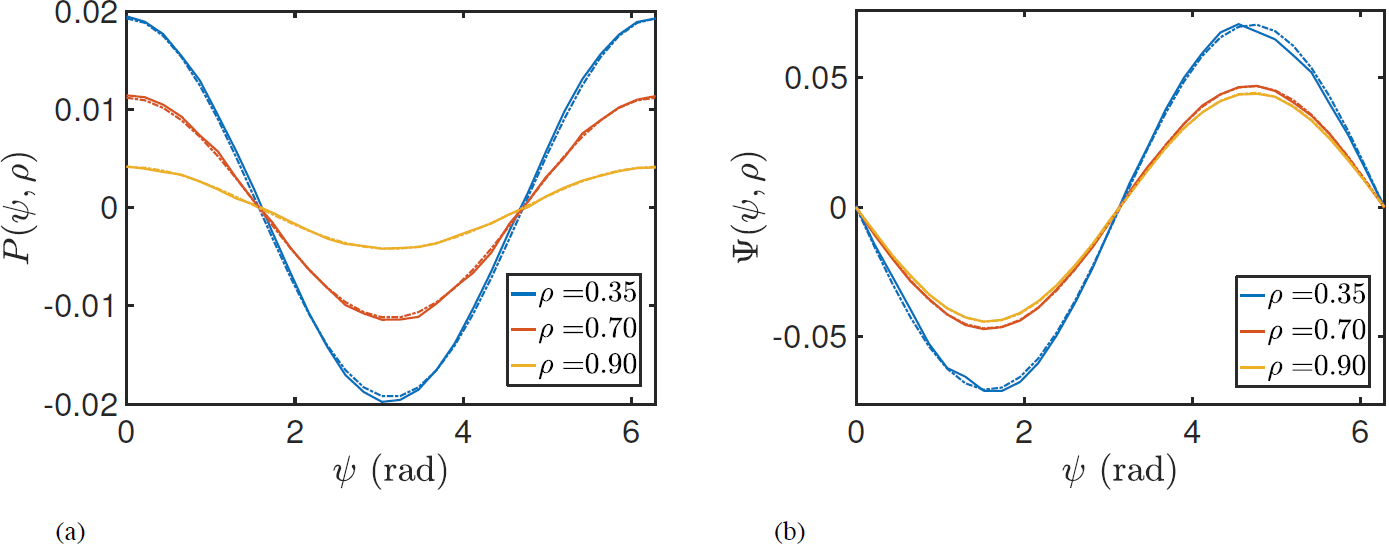
Instantaneous response curves as function of 0 for different values of synchrony *ρ*. Dashed lines were obtained from Eqs. (22) and (23). Panels (a) and (b) show the ARC and PRC, respectively. Solid lines were calculated by simulating a large ensemble of Kuramoto oscillators.

The functions *P*(*ρ*, *ψ*) and Ψ(*ρ*, *ψ*) are the instantaneous response curves for a population of oscillators with a phase response function of *Z*(*θ*) = — sin(6). This simplified case leads to very specific qualitative predictions. For both response curves, the effects of stimulation are predicted to be more magnified when stimulation is applied at lower amplitudes. In addition to this, whether or not stimulation has an amplifying or suppressing effect on the respective quantities is dependent only on the phase. In the absence of stimulation, the first term of Eq. (20) predicts the amplitude to decay with a rate proportional to the diversity of natural frequencies of individual oscillators 7 whilst the second term predicts the amplitude to increase according to a term proportional to the coupling strength k. Eq. (21) predicts the phase 0 to evolve according to w_0_.

To demonstrate the amplitude dependent effects of stimulation predicted by Eqs. (22) and (23) we simulate the Kuramoto model using a large number of oscillators (*N* = 3000). The stimulation amplitude *I* was chosen to be small at ≃ 0.04 and numerical integration was performed using the Euler method with a time step of δt ~ 0.001. For such a system and in the absence of stimulation, the magnitude of the order parameter *ρ* will tend asymptotically to [1 — (2/k)]^-0 5^ for *k* > 2 [29]. We can therefore fix the value of *ρ* in simulation by choosing an appropriate value of *k*. The parameters w_0_ and γ were not expected to affect the response curves and hence were arbitrarily chosen to be ω_0_ = 30 and 7 = 1. After the system has evolved to the asymptotic state, we provide stimulation at a particular phase over a single time step. The changes in *ρ* and 0 resulting from the perturbation divided by δt would then approximately equal to *P*(*ρ*, *ψ*) and *Ψ*(*ρ*, *ψ*), respectively. Figure 2 shows the response functions P(*ρ*, *ψ*) and *Ψ*(*ρ*, *ψ*) for different amplitudes *ρ* and also a comparison with results from simulating a population of Kuramoto oscillators.

Figure 2a and Eq. (22) shows the ARC is shifted with respect to *Z*(9) and that, for a given *ρ*, the most effective reduction of oscillation amplitude is achieved when phase-locked stimulation is provided at phase π. An intuition for these effects is shown in Figure 3a. The form of the phase response function *Z*(*θ*) leads to a region of phases for which the oscillators will either slow down or speed up upon stimulation. Stimulation applied to a population of oscillators corresponds to a perturbation with a differential effect across the system of oscillators, i.e. with some oscillators responding differently to others and depending on their phase. It is this differential effect which gives rise to changes in the width of the oscillator distribution and therefore changes in amplitude. In particular, if stimulation is applied when the distribution of oscillators is centred around 180 degrees, stimulation is shown to have a desynchronising effect as half the oscillators will speed up and the other half will slow down, as illustrated in Figure 3a.

An intuition for the amplitude dependent effects can also be seen in Figure 3a. If we consider the case where the system is strongly synchronised (orange curve), then stimulation can have little effect on amplitude since all oscillators would be perturbed by a similar amount. If we now increase the width of the oscillator distribution (red curve) and the amplitude reduces, then the differential effects gives rise to a greater amplitude change, thus stimulation at lower amplitudes leads to magnified effects. It is also evident from Figure 2 that there exists a relationship between the ARC and slope of the PRC. In particular, the ARC is negative at those phases for which the slope of the PRC is positive. One can also see from Eqs. (22) and (23) that for a given *ρ*, the ARC (P(*ρ*, *ψ*)) is proportional to the negative derivative of the PRC with respect to *ψ* 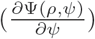. We will later analyse how this relationship generalizes.

**Fig.3.**
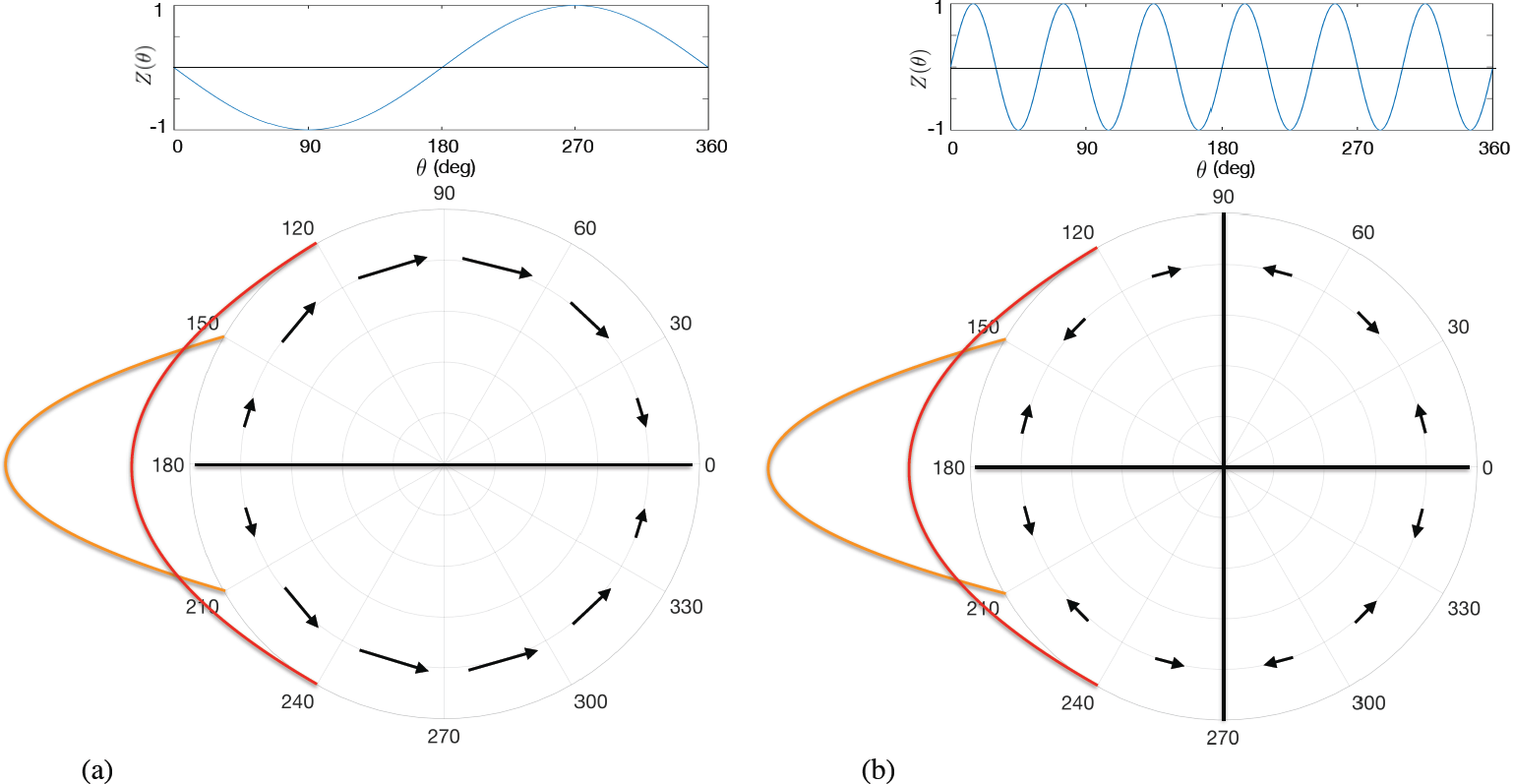
Intuition behind amplitude dependent effects on the ARC. For each panel, the top plot shows the phase response function for an individual oscillator *Z*(*θ*). The bottom part shows the shape of this function as represented by arrows indicating the effect of stimulation at a particular phase. The length of the arrows reflect the magnitude of phase change due to stimulation. The orange and red curves schematically show distributions of oscillators centred around *ψ* = 180°, as discussed in the text. (a) For the single harmonic case of *Z*(*θ*) = — sin(*θ*), the amplitude dependent effects are predicted to be monotonic, with magnified effects at lower amplitudes. (b) For the higher harmonic case of *Z*(*θ*) = sin(6*θ*) a non-monotonic relationship is predicted.

### Generalised response curves

In this subsection we consider the case where the phase response function *Z*(*θ*) has a general form, since the phase response curves of biological neurons may have diverse shapes [26]. We start by providing an intuition for why the qualitative effects of stimulation are expected to be different when *Z*(*θ*) contains higher harmonics. For the case of *Z*(*θ*) = sin(6*θ*), as shown in Figure 3b, and the simple case of oscillators distributed around 180 degrees, the effects of stimulation may actually be greater at larger amplitudes - which is in contrast to our results for the single harmonic case. In the high amplitude regime, as shown by the orange curve, the qualitative effects of stimulation are predicted to be similar to that of the single harmonic case. At lower amplitudes, some of the oscillators will be shifted away from the centre of the distribution while other towards it, reducing the overall effect of stimulation. To analyse this more formally, we now consider the case where *Z*(*θ*) takes the form of a general Fourier series

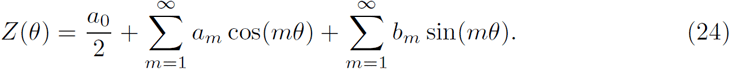

Using the results from Lai and Porter [30], an expression for the time evolution of the order parameter can be obtained,

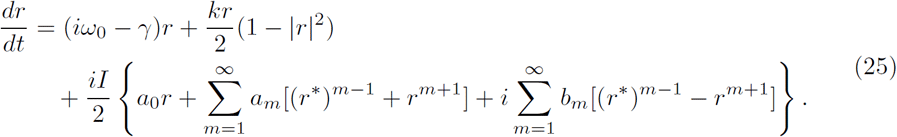

Inserting the expression for the order parameter (Eq. (7)) into Eq. (25), we find expressions for the time evolution of *ρ* and *ψ*, but now the instantaneous response curves for amplitude and phase, respectively, are given by (cf. [32])

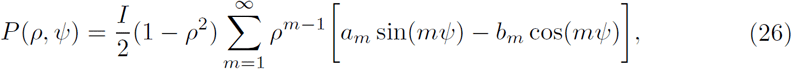

and

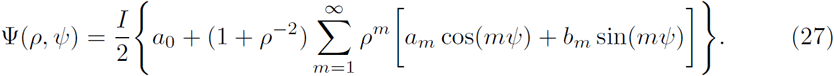

Eqs. (26) and (27) describe the instantaneous response curves for a system of oscillators whose distribution satisfies Eq. (16). Its worth noting that both equations are independent of the parameters of the Kuramoto model and are only dependent on the characteristics of an oscillator distribution satisfying the *ansatz.* It is known [30, 33, 34] that the presence of higher harmonic modes in the phase response function *Z* can cause the oscillators to cluster and lead to a breakdown of the *ansatz* given by Eq. (16). Lai *et al.* [30] investigated the effects of introducing noise through the phase response function and found good agreement between theory and simulation only when *Z* consisted of a dominant first harmonic mode. We therefore expect Eqs. (26) and (27) to reasonably approximate the response when *Z* has a dominant first harmonic and/or the stimulation amplitude *I* is small [35].

**Fig.4.**
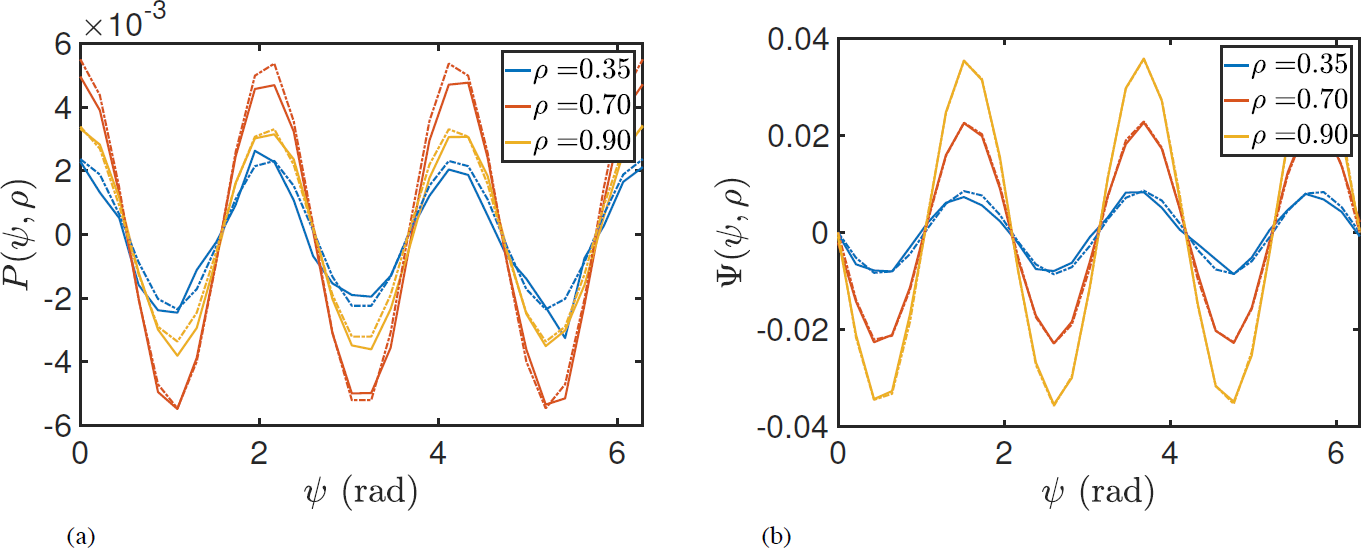
Instantaneous response curves as function of *ψ* for different values of synchrony ρ, for sample response function of individual oscillators including higher harmonic *Z*(*θ*) = — sin(3*θ*). Dashed lines were calculated from Eqs. (26) and (27). Panels (a) and (b) show the ARC and PRC, respectively. Solid lines show results from simulating a large ensemble of Kuramoto oscillators. In panel (b) theoretical predictions and simulations overlap so closely for some *ρ* that only a single curve can be seen.

Using the methodologies from before, we simulated a population of Kuramoto oscillators to demonstrate the predicted amplitude dependence of stimulation for the case of *Z*(*θ*) containing higher harmonics. Figure 4 shows an example of the ARC and PRC for the case of *Z*(*θ*) = — sin(3*θ*). In contrast to Figure 2, the effects of stimulation are not necessarily monotonic functions of *ρ*. This can be seen for the case of the ARC shown in Figure 4a, where the effects are magnified between *ρ* = 0.35 and *ρ* = 0.70 but are reduced between *ρ* = 0.70 and *ρ* = 0.90. For the PRC in Figure 4b, it is clear that the effects are monotonic but that the effects of stimulation now reduce with reducing amplitude, in contrast with the single harmonic case.

The expressions for the instantaneous response curves can be used to make qualitative predictions about how a subject should respond to stimulation. Eqs. (26) and (27) involve an expansion according to the harmonics of *Z*, with each term being the product of both a phase dependent part and an amplitude dependent part. If we restrict our analysis to the simple case where *Z* is well-approximated to be a single harmonic mode, then it can be seen that whether stimulation has a suppressive or amplifying effect on amplitude is dependent only on the phase and the magnitude of these effects is determined by the amplitude. Plots of the amplitude dependent part for several harmonic modes can be seen in Figure 5 for both the PRC and ARC. For the case of *Z* containing a single harmonic, each curve describes how the magnitude of the response curves is expected to change as a function of amplitude. Figure 5a shows the magnitude of the ARC as a function of *ρ* for several harmonic modes. A non-monotonic relationship is predicted only for higher modes *m* > 1, with the maxima occurring for each curve at

**Fig.5.**
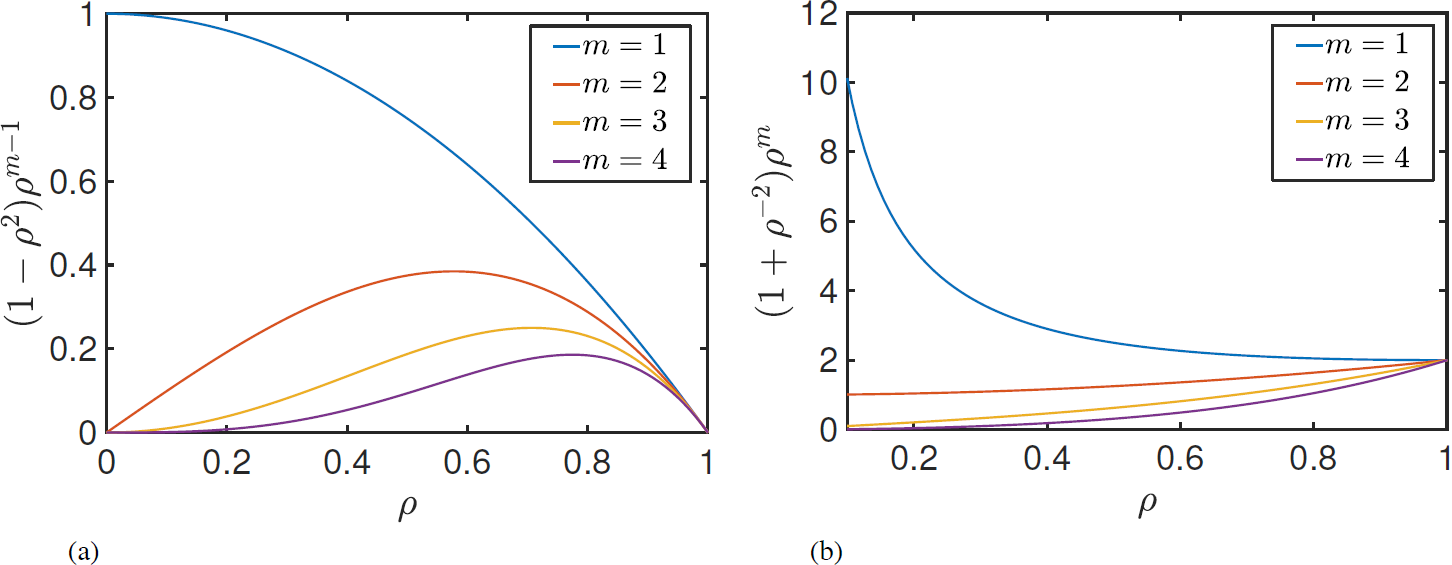
Plots of the amplitude dependent multipliers for each Fourier mode of the ARC (a) and the PRC (b). For a given *Z*(*θ*) with a single mth Fourier mode, the multipliers show how the size of the effects of stimulation are expected to change as a function of *ρ*.

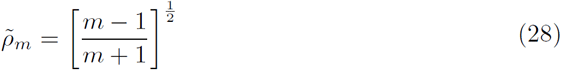

In the high synchrony regime 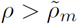, the gradient in each case is found to be negative, implying that when stimulation is applied at lower *ρ*, the effects of stimulation increase. In the low synchrony regime 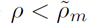, stimulation applied at lower amplitudes *ρ* is predicted to lead to smaller effects. For the case of the first harmonic mode, where 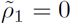 implies that the gradient is negative across the range 0 ≤ *ρ* ≤ 1, the predicted effects are particularly noteworthy as they are both quantitatively and qualitatively different from the other modes. Concisely, for *Z* consisting of a single first harmonic mode, we predict that delivering stimulation at lower *ρ* will result in greater effects, for all values of *ρ*.

Figure 5b shows the magnitude of the PRC as a function of amplitude for several harmonic modes. Here we find the qualitative differences between the effects of the first harmonic mode and higher modes to be even more notable. A monotonic relationship is predicted across the harmonic modes but with differing gradients between the first and higher modes. The condition for positive (or zero) gradients is that 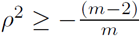, which is only satisfied for *m* > 1, hence for higher modes, a positive (or zero) gradient can be found for all *ρ* in contrast to a negative gradient for all *ρ* for the first mode. Qualitatively, this means that for Z consisting of a dominant higher harmonic, delivering stimulation at lower *ρ* will result in smaller changes in phase. For *Z* consisting of a dominant first harmonic, the opposite is predicted, namely delivering stimulation at lower *ρ* will result in larger effects. These effects become particularly apparent if the oscillation amplitude is close to 0, where the ρ^-1^ term attached to the first harmonic begins to become very large.

Finally, it is worth noting that for stimulation to have an effect on amplitude, the function *Z*(*θ*) does not need to have a region where *Z*(*θ*) < 0 (i.e. it does not need to be of type II). Although in the examples we used *Z*(*θ*) with negative regions, *Z*(*θ*) can be shifted by adding a constant a_0_, and this constant will not affect ARC, because it does not appear in Eq. 26. A critical condition for the stimulation to have an effect on amplitude is that the function *Z*(*θ*) is not constant, which allows the stimulation to have a differential effect on oscillators in different phases.

### Relationship between averaged response curves

Existing experimental studies have reported the ARC and PRC averaged across amplitudes of pathological oscillations. In this subsection, we study the properties of such averaged curves. In the next subsection we use this relationship to test if the response to stimulation of individual patients is well described by the Kuramoto model.

Expressions for the averaged response as a function of *ψ* can be obtained by taking expectation values using the function *h(*ρ**|*ψ*), which is the probability density function for the system being in a state *ρ* given a phase *ψ*. In the absence of stimulation, the dynamics of *ρ* for the Kuramoto system are phase-shift invariant. Therefore, if the effects of stimulation are small, it is reasonable to assume that *h(*ρ*|*ψ*) ≃ h(p)* and

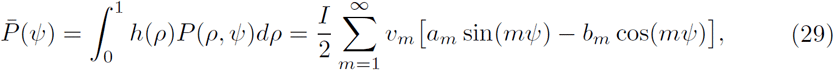

and

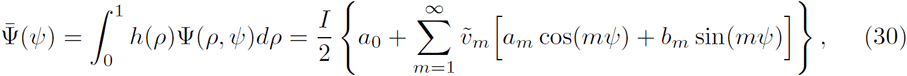

where

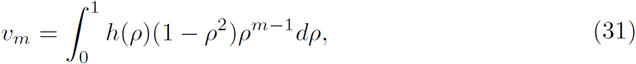

and

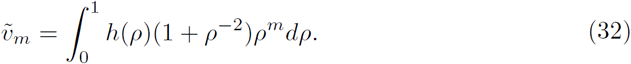

We first describe the relationship between the ARC and PRC for the cases where *Z* contains a single dominant harmonic as in these cases clear predictions can be made by the model. First, we use the derivative of Ψ(*ψ*) giving

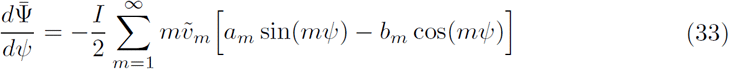

and also by dividing *P*(*ψ*) by 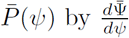 leads to

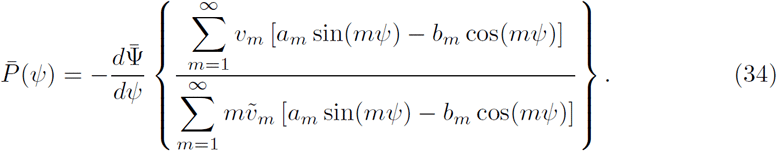

Now considering the case where *Z* contains only the *q*th harmonic

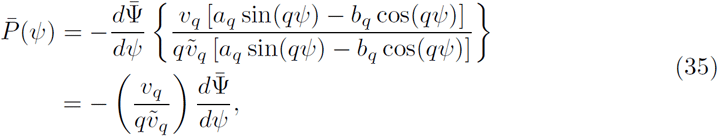

which shows that in the cases where Z contains a single harmonic the averaged ARC is a scaled version of the negative gradient of the averaged PRC.

Since the constants *V*_q_, 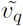 and *q* are all positive, it is expected that for *Z* containing a single or dominant harmonic, a strong positive correlation should exist between 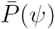 and 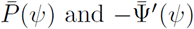. To investigate the correlation coefficient for higher harmonics, we simulated the Kuramoto model using randomly generated phase response functions. The parameters of the Kuramoto model (*N*, *ω*_0_, γ) were the same as in previous simulations. The number of harmonics *(n*_h_) to use in each function was chosen sequentially from 1 to 4. In each case, the harmonics of each function were chosen to be a random subset of size n_h_ from the set {1, 2, 3, 4}. A random phase response function *Z* was generated by choosing a set of coefficients {*a*_m_} and {b_m_} (which includes randomising a_0_) whose values were sampled from a standard normal distribution. For each *Z* with a given number of harmonics, the response curves as a function of phase were calculated at values of synchrony *ρ* = {0.4,0.6,0.8}. The phases were chosen from a uniformly spaced grid between 0 and 2n. The response of the system was taken to be from a single pulse of stimulation. For a given number of harmonics, 30 averaged response curves were calculated. The correlation coefficient between 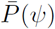 and 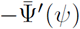 was then calculated by first averaging *P*(**ρ**, **ψ**) an *Ψ*(**ρ**, 0) across *ρ* and then calculating 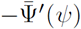 by averaging the gradient in the forward and backward direction around a particular phase *ψ*. Figure 6 shows the value of the correlation coefficient to be only slightly affected by increases in the number of harmonics. This implies that if a system is well described by the Kuramoto model, then a strong positive correlation between 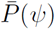 and 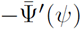 should exist.

**Fig.6.**
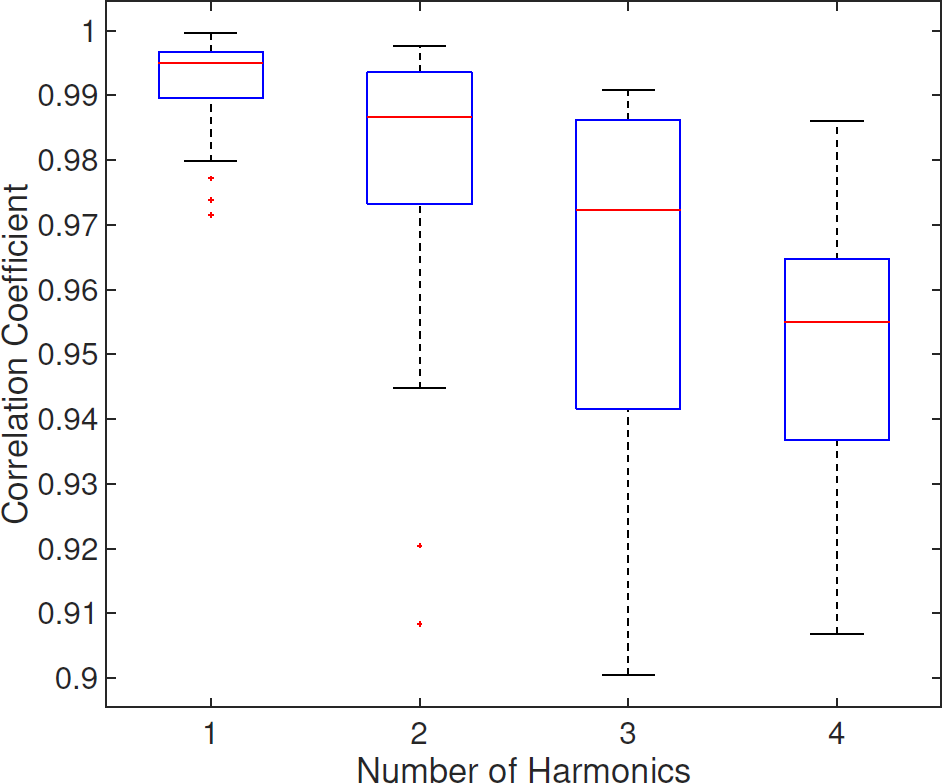
Box plots showing the correlation coefficients between P(*ψ*) and—f’(0) calculated by generating a random phase response functions Z with differing number of harmonics.

The results of this subsection extend previous observations [21, 36] that stimulation is most effective when applied at the phase which maximizes Z’(*θ*). However, the phase response curves of individual oscillators *Z*(*θ*) are difficult to measure. Nevertheless, Eq. (35) shows an analogous relationship to exist between averaged population response curves. Together with Eq. (11), the above analysis predicts that there also should exist a positive correlation between experimentally measured ARC and the gradient of experimentally measured PRC.

### Comparison with experimental data

The results presented in previous sections provide a framework within which we can make some qualitative predictions about how a patient should respond to phase-locked DBS. In this subsection we compare two key predictions of the theory with experimental data:

1. There should exist a strong correlation between averaged ARC 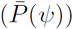 and the negative gradient of the averaged PRC 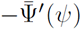.
2. If the phase response function *Z* contains a dominant first harmonic, then the effects of stimulation should be magnified if it is applied when the amplitude of oscillation *ρ* is low.

We tested these predictions using data from the study of Cagnan et *al.* [16]. In this study, phase-locked DBS was delivered according to the tremor measured by an accelerometer attached to the patient’s hand. Data was collected from 6 ET patients and 3 dystonic tremor patients. We investigated the 5 ET patients that exhibited a significant response to stimulation. The data from these 5 patients was associated with 6 hemispheres, with datasets 4L and 4R denoting tremor data for the left and right hand of Patient 4, with stimulation delivered to the contralateral hemisphere.

The tremor data was filtered using a Butterworth filter of order 2 with cut-off frequencies at ±2 Hz around the tremor frequency. Stimulation was delivered over a set of trials (typically 9), with each trial consisting of 12 blocks of 5 second phase-locked stimulation at a randomly chosen phase from a set of 12. Each block of phase-locked stimulation was also separated by a 1 second interblock of no stimulation. The envelope amplitude and instantaneous phase were calculated using the Hilbert transform. The averaged amplitude response for a particular phase was calculated to be the difference between the average envelope amplitude within a 1 second window before the end of the stimulation block and the average envelope amplitude within a 1 second window prior to the onset of stimulation. The averaged phase response was calculated using a similar methodology. The unwrapped phase was calculated for the data 1 second prior to the onset of stimulation. A linear function was fitted to this phase evolution and extrapolated to the end of the stimulation block. The value for the instantaneous phase obtained using this extrapolation could be taken as the expected phase of the system in the absence of stimulation. The difference between the actual phase at the end of the stimulation block and this expected phase was taken to be the phase response. In both cases, the responses for a particular phase were averaged over all trials in the dataset. The derivative of the PRC with respect to *ψ* was calculated numerically by averaging the gradient in the forward and backward direction around a particular phase *ψ*. To determine the effects of amplitude on the response curves, we use a ‘single pulse’ method, where the data is binned at low, medium and high amplitudes, with each bin containing the same number of points. Within each bin we calculate the response of the system from a single pulse of stimulation. In the case of the amplitude response, we calculate the difference in the mean of the amplitude after and before the pulse. The data used for calculating the mean in each case is taken between pulses. In the case of the phase response, a straight line is fitted to the unwrapped phase evolution from before the pulse. The phase response is taken to be the difference between the actual phase and the linear extrapolation evaluated at a point after the pulse and just before the next pulse.

We first tested Prediction 1 for each patient and excluded those patients which either do not have significant correlation *(p* < 0.05) or have negative correlation. The lack of significant correlation between 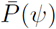 and 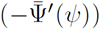 could indicate that the patient is not well described by the Kuramoto model or that the response curves have not been accurately determined. We therefore restrict subsequent testing of Prediction 2 to only those patients who exhibit significant correlation. To infer whether the phase response function *Z* for a given patient contains a dominant first harmonic, we used the property illustrated in Figure 5b, namely that the magnitude of the PRC should increase with reducing *ρ* only if *Z* contains a single first harmonic. For such patients, we would then expect the magnitude of the ARC to increase with reducing *ρ*. Table 1 gives the correlation coefficients between 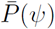 and 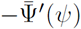 for each patient where DBS was found to have a significant effect on tremor. From these, significant correlation was only found for Patient 5, with the correlation coefficient being (0.72). The response curves for this patient are shown in Figure 7. Table 1 also lists uncorrected p-values for the correlation. Since we analysed data from 6 datasets, the Bonferoni corrected p-value of correlation for Patient 5 would still be significant (*p* = 0.048). In summary, we did not find strong support for Prediction 1, as the correlation between the ARC and the negative gradient of the PRC was significant in only 1 out of the 6 datasets analysed. As outlined above, we then tested Prediction 2 only for Patient 5 who fulfilled Prediction 1. To determine the effects of amplitude on the response curves, we analysed the response of the system using the single pulse method. This is because the amplitude of tremor can vary substantially within the 5-second stimulation intervals. Figure 8 shows the response curves at 3 amplitude bins for Patient 5. To quantify the magnification of the response curves, we compute the standard deviation across the phase bins. The uncertainty on the standard deviation *σ* is calculated using the method of propagating errors [37]. We find this to be most appropriate here as it allows us to incorporate the errors of the binned response curves. For a dataset consisting of *M* points *Y* = {*yj*} with mean *ȳ* and corresponding standard errors *ΔY* = {Δ*y*_*i*_}, we find the error on the standard deviation *Δσ* to be

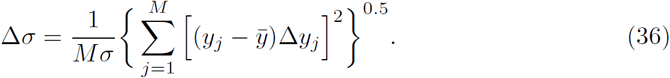

Figure 9b shows the extent of magnification across 3 amplitude bins for the PRC. For Patient 5, we find evidence for increasing magnification with reducing amplitude, which when taken together with the positive correlation between 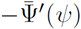 and 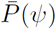, is indicative of a Kuramoto system with a phase response function *Z* containing a dominant first harmonic mode. For such a system, Prediction 2 states that we should also find increasing magnification of the ARC with reducing amplitude. Figure 9a shows the extent of magnification across 3 amplitude bins for the ARC. Here we can see evidence for increasing magnification with reducing amplitude for Patient 5, which agrees with our predictions and was the only patient to exhibit such effects across all the amplitude bins. For completeness, Figure 9 also shows the results of the above analysis for other patients. It is evident in Figure 9b that, for all patients, the magnification of the PRC is higher for stimulation at lower ρ. Interestingly, Figure 9a shows that the magnification of the ARC for stimulation at lower *ρ* is only exhibited by Patient 5, which is also the only patient who had significant correlation between the ARC and negative gradient of the PRC.

**Table 1.**
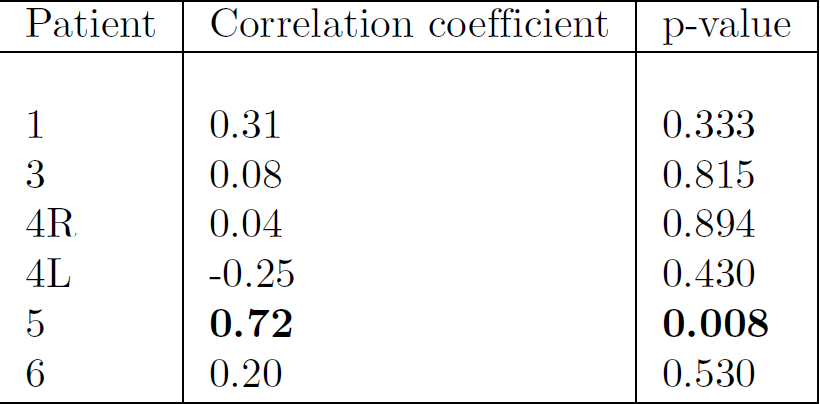
Table showing the correlation coefficients and corresponding p-values between *P̄*(*ψ*) and —*Ψ*’(*ψ*) for each ET patient with a significant effect of stimulation on amplitude of tremor, reported in the study of Cagnan *et al.* [16].

**Fig.7.**
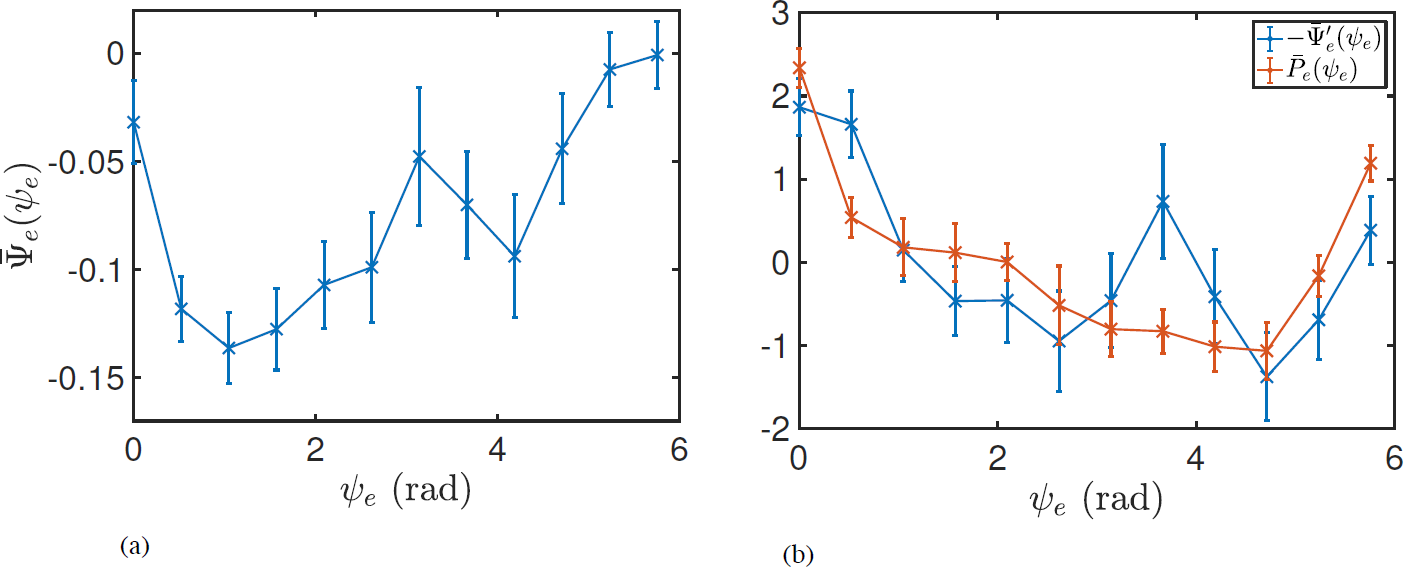
Plots showing the measured ARC and the negative gradient of the PRC for Patient 5. The curves were z-scored to have comparable scale. The error bars indicate the standard error of the mean.

**Fig.8.**
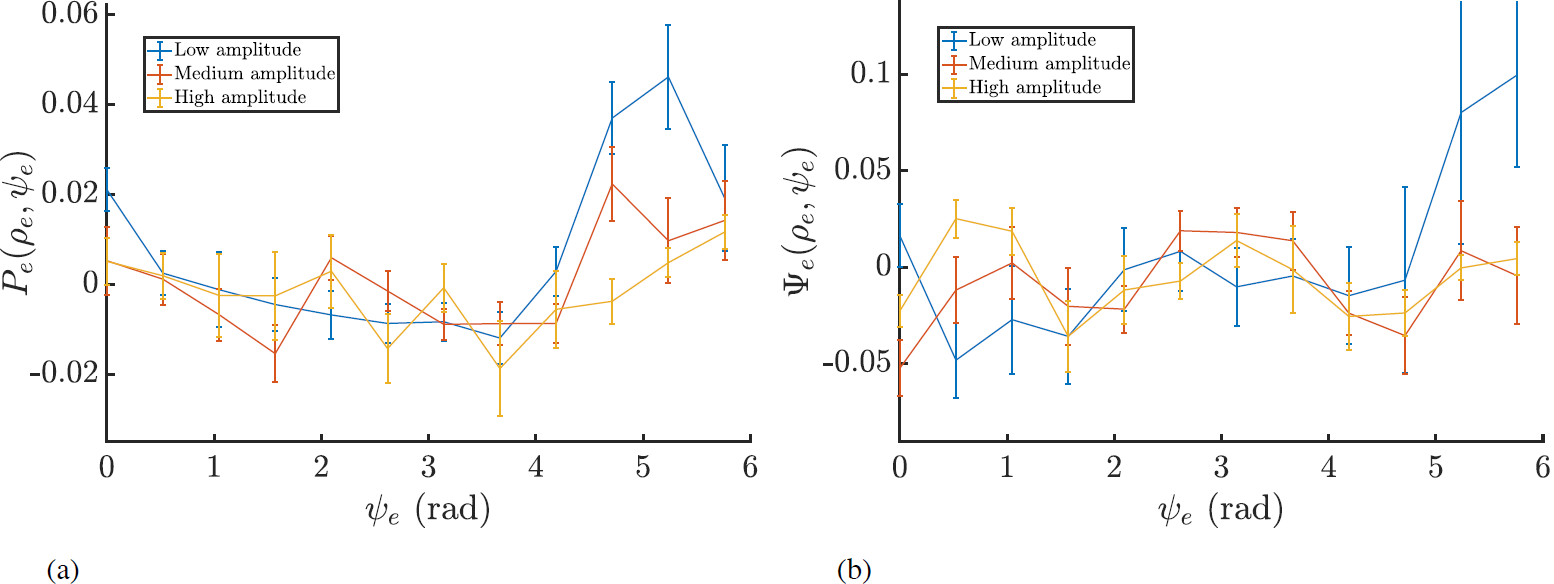
Response curves as a function of for Patient 5 calculated by binning according to amplitude. Panel (a) shows the ARC and (b) shows the PRC. The error bars are the standard error of the mean.

**Fig.9.**
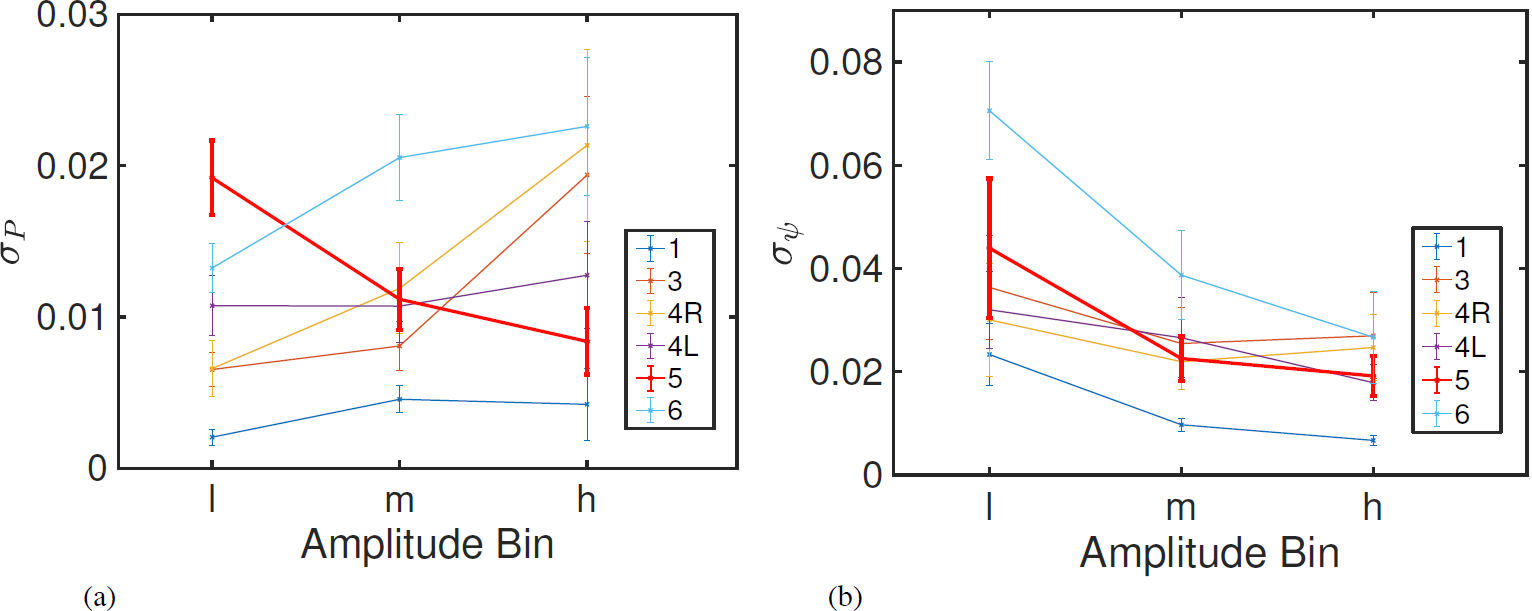
Magnification of the response curves calculated using the standard deviation for 3 amplitude bins low (l), medium (m) and high (h). Panel (a) shows the magnification for the ARC and (b) shows the magnification for the PRC.

## Discussion

We have presented a framework for testing the theory that oscillations found in tremor data can be represented by a Kuramoto system, whose behaviour is described by Eq. (13). The theories we present make clear qualitative predictions about the amplitude dependence of the response curves which we can test using experimental data. The effects of stimulation on tremor amplitude are summarized in Figure 10. For the cases where the phase response function *Z*(9) contains a single dominant harmonic, whether the effects of stimulation are suppressive or amplifying depends on the phase at which the stimulation is applied (compare columns), while the magnitude of the stimulation effect depends on the amplitude of oscillations (compare rows) at the point of stimulation. If the phase response function has a dominant first harmonic, the effect is largest when the tremor amplitude is lower (illustrated in the figure), while for phase response functions with a dominant higher harmonic, the magnitude of the effect is a non-monotonic function of the amplitude. The magnitude of the phase shift of the signal due to stimulation also depends on the phase response function. If it has a dominant first harmonic, then the effects should increase with reducing amplitudes; thus the oscillation should get most easily entrained to phase-locked stimulation if the amplitude of the oscillation is small. For phase response functions containing a dominant higher harmonic, the effects of stimulation should increase with increasing amplitude. The analysis of population response curves in this paper is similar to that presented by Hannay et al. [32]. The formula for the PRC given in Eq. (27) is the same as that given by Hannay et al. They also derive a similar expression for the ARC, but define the ARC to be the ratio of the amplitude post and pre stimulation, while we define it to be the difference between these amplitudes, which is closer to the convention used in the DBS literature. Here we extend their analysis of response curves in a way which is more relevant for designing adaptive DBS.

**Fig.10.**
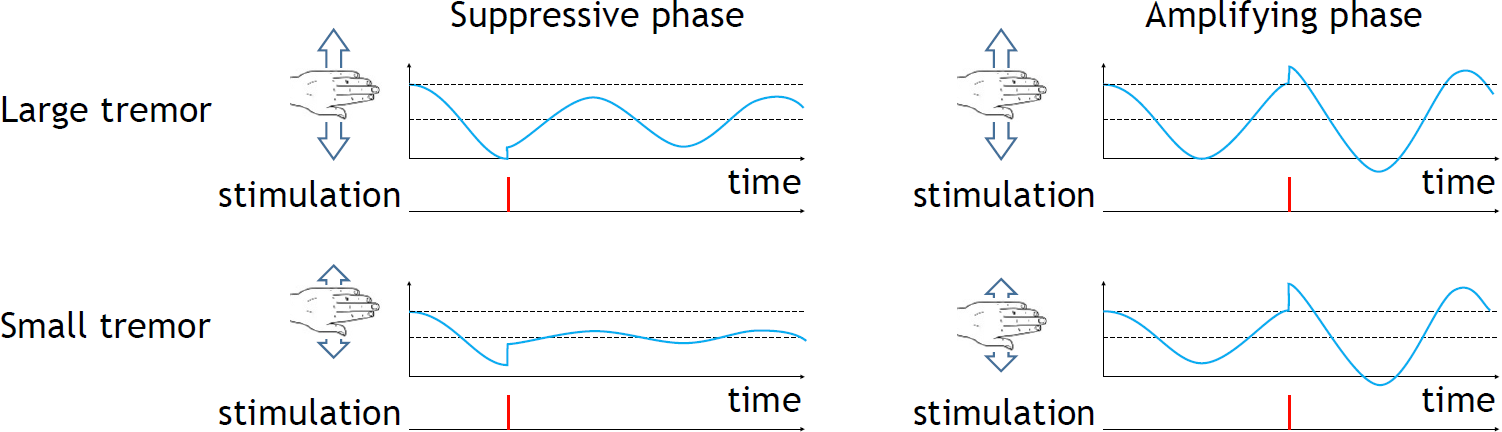
Schematic illustration of the effects of stimulation at different phases (columns) and different amplitudes (rows), when the phase response function of individual oscillators has a dominant first harmonic. In each display, blue curve shows the tremor signal, dotted lines indicate the maximum and mean of the signal before stimulation, and red bar indicates the time of stimulation.

### Relationship to experimental data

We find good agreement between our theories and the data from one patient, namely a strong positive correlation between 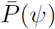 and 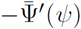 together with increasing magnification of the PRC with reducing amplitude is associated with an increasing magnification of the ARC with reducing amplitude.

The lack of significant correlation between 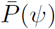 and 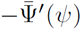 for the other patients could be due to experimental noise, which may prevent the response curves from being determined accurately. However, another possibility is that these patients are simply not well described by the models presented here. This could be due to the various assumptions, both about the nature of stimulation and the form for the distribution of oscillators, which are used to derive the theoretical response curves. Since the theoretical response curves are only valid for those distributions satisfying the *ansatz*given by Eq. (16), the response curves for any distribution deviating from this form are expected to be different from those predicted. Distributions not represented by the *ansatz* include those clustered configurations which can arise through random effects or stimulation applied through phase response functions containing higher harmonic modes. Eq. (13) also assumes that each oscillator responds only according to a single phase response function *Z*(9), which may not be a good approximation for certain patients. Since we interpret an oscillator as representing the activity of neurons or micro-circuits, it follows that these neurons should have some spatial separation in the brain [38] and hence experience stimulation differently depending on their location relative to the electrode. This is not an effect which is captured by the models presented here. Furthermore, our computational model does not capture the effects of synaptic plasticity triggered by stimulation. Presence of such plasticity may be suggested by a delayed appearance of tremor following offset of the prolonged phase-locked stimulation [16]. Finally, ET is known to be a heterogeneous disorder [39] and different patients are likely to have different underlying pathologies. As a result of this, the assumptions used in the model may need to differ depending on the patient.

### Hybrid DBS

In light of our predictions, we propose a new strategy for DBS which may be effective for certain patients, namely those for whom the effects of DBS are magnified when stimulation is applied at lower amplitudes. The approach, illustrated in Figure 11, combines the aforementioned phase-locked and adaptive approaches described in Figure 1. The general idea is to only apply high frequency DBS at high amplitudes in order to drive the tremor into low synchrony regimes that are more susceptible to phase-locked DBS. With such a control strategy, the high frequency DBS would be used less often than in the current adaptive DBS approach, as it would only be used to bring the tremor into the low synchrony regime, at which point phase-locked DBS would suppress the amplitude further, thereby keeping the system in this mode.

Separating the high and low synchrony regimes can be done using an amplitude threshold, in a similar way to the adaptive DBS approach [15]. When the amplitude of oscillations exceeds the threshold, then high frequency DBS would be delivered since the patient would be in the high synchrony regime. If the amplitude of oscillations falls below the threshold, then phase-locked DBS is delivered, since the patient would be in the low synchrony regime. In addition to the various parameters of stimulation, the choice of threshold is also likely to be an important factor in determining the overall efficacy of the method [40].

**Fig.11.**
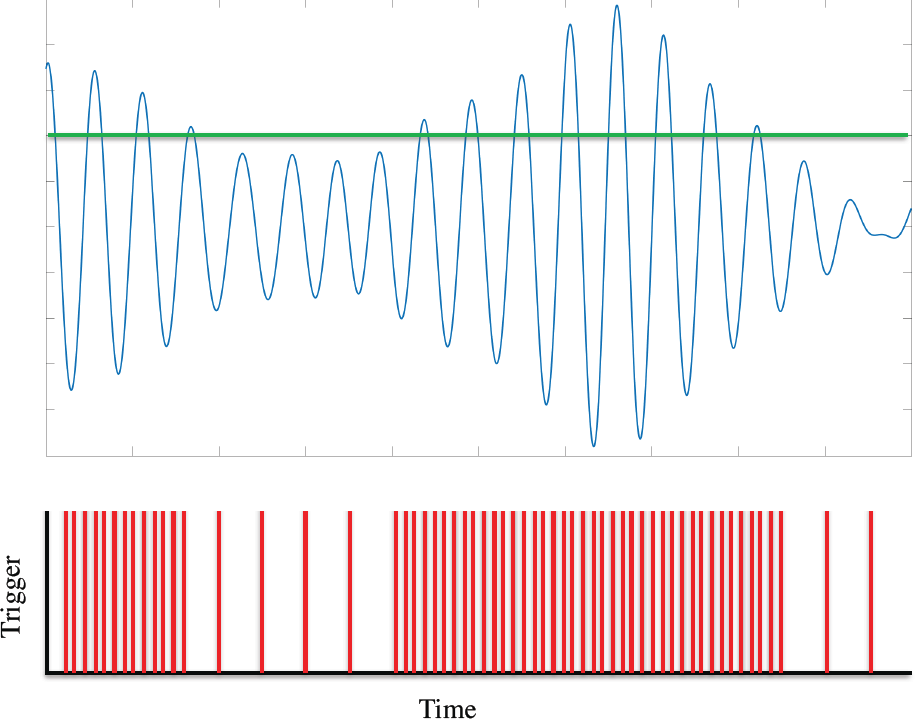
Hybrid DBS strategy. High frequency DBS is applied when the amplitude of oscillations exceed a predefined threshold. Below this threshold, phase-locked DBS is applied.

### Future Work

The theories we have presented here leave plenty of scope for future work. On the theoretical side, investigating the effects of clustering on the response curves in addition to considering a multi-population Kuramoto model would be two possible avenues to explore. Its also worth mentioning that the instantaneous response curves given by Eq. (26) and Eq. (27) are independent of the parameters of the Kuramoto model and can be derived using only the assumption of the Ott and Antonsen *ansatz.* Therefore, in principle, they should be valid for other systems for which the Ott and Antonsen *ansatz*can be applied, such as an infinite network of theta neurons [41]. Kuramoto-like phase models arise through the phase reduction of oscillating units. By contrast, theta neurons can be in both excitable and oscillatory regimes while still being amenable to the Ott-Antonsen reduction [42]. It would be interesting to relate the phase-response curves of the population dynamics [43] to the data presented here. Furthermore, the model analysed in this paper assumes a Cauchy distribution for the natural frequencies, which has very long tails, and other distributions (e.g. Gaussian) may be a more realistic description of neuronal frequencies. It has recently been demonstrated that the Ott and Antonsen *ansatz* can be applied to systems where the natural frequencies are Gaussian distributed [44], and it would be interesting to extend the analysis of population response curves to this case.

On the experimental side, there is the question of the hybrid DBS strategy, whose efficacy remains to be determined. To further strengthen our conclusions, we also hope to perform our analyses on more data. Given that testing our theories is conditional on finding patients for whom P(*ψ*) is correlated to — T’(−0), a study such as ours would greatly benefit from a larger dataset, both in terms of the number of patients and the length of time each patient is stimulated for. The latter would allow us to determine the response curves more accurately, which is expected to be particularly beneficial for our methods. Obtaining longer datasets poses a challenge due to the inherent difficulties associated with recording from patients, particularly the onset of fatigue. Alternatively, improvements to the accuracy of the response curves could be realised by improving the methods used to calculate them.

## Footnotes

1 Note that Eq. (17) assumes that individual neurons react to stimulation in the same way as to input from other neurons.

